# PRODIGY: personalized prioritization of driver genes

**DOI:** 10.1101/456723

**Authors:** Gal Dinstag, Ron Shamir

## Abstract

**Background:** Evolution of cancer is driven by few somatic mutations that disrupt cellular processes, causing abnormal proliferation and tumor development, while most somatic mutations have no impact on progression. Distinguishing those mutated genes that drive tumorigenesis in a patient is a primary goal in cancer therapy: Knowledge of these genes and the pathways on which they operate can illuminate disease mechanisms and indicate potential therapies and drug targets. Current research focuses mainly on cohort-level driver gene identification, but patient-specific driver gene identification remains a challenge.

**Methods:** We developed a new algorithm for patient-specific ranking of driver genes. The algorithm, called PRODIGY, analyzes the expression and mutation profiles of the patient along with data on known pathways and protein-protein interactions. Prodigy quantifies the impact of each mutated gene on every deregulated pathway using the prize collecting Steiner tree model. Mutated genes are ranked by their aggregated impact on all deregulated pathways.

**Results:** In testing on five TCGA cancer cohorts spanning >2500 patients and comparison to validated driver genes, Prodigy outperformed extant methods and ranking based on network centrality measures. Our results pinpoint the pleiotropic effect of driver genes and show that Prodigy is capable of identifying even very rare drivers. Hence, Prodigy takes a step further towards personalized medicine and treatment.

**Availability:** The Prodigy R package is available at: https://github.com/Shamir-Lab/PRODIGY.

## Introduction

Cancer is an evolutionary process in which normal cells accumulate genomic and epigenomic alterations of various kinds including Single Nucleotide Variations (SNVs) and chromosomal aberrations. Some of these alterations confer growth and positive selection advantage to the mutated cells, giving rise to intensive proliferation and tumors (Stratton *et al.*, 2009). The alterations can be inherited through germ line mutations as in the case of *BRCA1* and *BRCA2* in breast cancer(King *et al.*, 2003) or occur somatically (Stratton *et al.*, 2009). While somatic mutations also occur in normal cells, they are neutral or cause apoptosis but do not lead to transformation into a cancer cell.

### Driver mutations

Mutational events that grant selective growth advantage to the cell and thus “drive” it into tumorigenesis are called *driver mutations* (or driver events), and the genes in which these mutations take place are called *driver genes*. In contrast, other mutations that occur in the same genome with the driver mutation but have not effect on fitness are called *passenger mutations* (Vogelstein *et al.*, 2013).

The overall number of observed mutations varies among tumor tissues. Kim & Kim (Kim and Kim, 2018) analyzed dozens of cancer patient cohorts from TCGA (Weinstein *et al.*, 2013) and found that the average number of somatic mutations can reach up to thousands per tumor in some cancer subtypes. There is a very extensive debate regarding the number of driver mutations among the observed mutations in each tumor (Stratton *et al.*, 2009; Vogelstein *et al.*, 2013; Anna C. Schinzel, 2008) but the consensus is that this number is very low. Obviously, there are many factors that contribute to the variation in the number of drivers, including the progression stage of the tumor (Vogelstein and Kinzler, 2015), its tissue of origin (Kandoth *et al.*, 2013), environmental properties such as smoking (Govindan *et al.*, 2012) and other factors like age (Xie *et al.*, 2014). Tomasetti et al. (Tomasetti *et al.*, 2015) showed that as little as three driver mutations suffice to develop lung and colorectal cancer. Nordling (Nordling, 1953) and Armitage (Armitage and Doll, 1954) suggested six or seven as the typical number of drivers.

It is therefore a challenge to distinguish driver from passenger mutations. The need to do so has high priority in cancer research - and in personalized cancer medicine in particular - for several reasons: 1) knowledge of the drivers and the mechanisms by which they operate can suggest potential treatments and drug targets. 2) Basing cancer treatment on molecular signatures rather than on the disease organ offers the opportunity to treat individuals with regimens not yet considered for their specific type of cancer. For example, many “basket” clinical trials, in which a specific drug is given to patients with diverse cancer types based on specific biomarkers, show that the same drug can have high efficiency across different types if the right mutation is detected (Hyman *et al.*, 2017).

### Driver gene identification in large cohorts

Computational research regarding driver genes first focused on distinguishing driver mutations from passengers in a cohort of patients (usually of the same tissue of origin): MuSiC (Dees *et al.*, 2012) uses the statistical significance of higher than expected rate of mutations, along with pathway mutation rate and correlation with clinical features to detect drivers. MutSigCV (Lawrence *et al.*, 2013) estimates the background mutation rate of each gene and identifies mutations that significantly deviate from that rate. MEMo (Ciriello *et al.*, 2012) tries to find small subnetworks of genes that belong to the same pathway and exhibit internal mutual exclusivity patterns. HotNet2 (Leiserson *et al.*, 2014) incorporates knowledge from protein-protein interaction (PPI) networks to find small subnetworks of frequently mutated genes using heat-diffusion process. TieDie (Paull *et al.*, 2013) also incorporates PPIs and mRNA expression data to find overlapping subnetworks that possess high degree of mutation and expression values using heat-diffusion. DriverNet (Bashashati *et al.*, 2012) tries to find a parsimonious set of mutated genes that is linked to genes that experience deregulation of mRNA expression in a given PPI network. Paradigm-Shift (Ng *et al.*, 2012) utilizes SNV, copy number variation (CNV), expression and known pathways to infer gain or loss-of-function of mutated genes in single patients. Many more methods for driver gene detection in cohorts are covered by Chang et al. (Cheng *et al.*, 2016) and Tokahim et al. (Tokheim *et al.*, 2016).

The methods above focus on general driver gene detection, but do not aim to offer personalized means of diagnosis or treatment: individual patients may have different compositions of mutated driver genes (**Supp. Fig. 1**). In addition, these methods rely on statistical power obtained by large cohorts and by doing so, they inevitably underestimate the importance of rare drivers that occur in only a handful of patients (also known as the “long tail phenomenon” (Garraway and Lander, 2013)) and are important only for them. Here we focused on patient specific driver prioritization.

Although many driver mutations were experimentally validated (Futreal *et al.*, 2004), personalized driver prioritization is needed for several reasons: 1) Some patients carry mutations in dozens of known drivers (**SFig 1**), and it is essential to understand which are the true drivers for the patient. 2) Some patients do not possess mutations in any known driver (**SFig 1**), so one has to find putative ones de novo. 3) Even if a patient has only few mutations in known drivers, and assuming they are all active, we still need to internally rank them, since the number of therapies that can be given to an individual at the same time is very low due to toxicity and adverse events (Kroschinsky *et al.*, 2017; Park *et al.*, 2013).

### Personalized driver gene profiles

To address the need for personalized driver gene identification and prioritization, one must develop methods that can operate on the data of a single patient. Several attempts have been made in this direction: DawnRank (Hou and Ma, 2014) uses a variant of Google’s PageRank to rank an individual’s mutated genes profile according to its effect on expression deregulation of downstream genes in a large directed PPI network. It ranks the genes by quantifying the impact of each of them on the differentially expressed genes (DEGs) using a diffusion process. SCS(Guo *et al.*, 2018) tries to find a parsimonious set of mutated genes that are linked to downstream DEGs in a large directed PPI network. These methods rank putative driver genes for a patient. In contrast, Hitn’DRIVE (Shrestha *et al.*, 2017, 2014) outputs a set of candidate driver genes without internal ranking. It tries to find a parsimonious set of mutations with short expected path lengths to a set of DEGs. The lack of ranking is a drawback from a treatment perspective, especially when the number of predicted genes is large.

### This study

Here we develop a new algorithm for ranking of driver genes of an individual. The algorithm, called PRODIGY (Personalized Ranking Of DrIver Genes analYsis) scores mutations by their influence on deregulation of multiple known pathways. Unlike the methods described above, Prodigy collects multiple signals from many local views of the same tumor rather than one global view. These local views are based on curated pathways and each one reflects a different aspect of the deregulation state of the tumor. Thus, the extent to which a given mutation explains multiple pathway deregulations serves as a proxy to the likelihood that this mutation is indeed one of the drivers. Our algorithm assumes that driver mutations influence the deregulation of other genes in affected pathways. In particular the true drivers will have good connectivity to these pathways, and our method is designed to score such connections correctly using a variant of the prize collecting Steiner tree problem (PCST). By aggregating many local views for all mutations of an individual, a global picture is obtained and the personalized landscape of drivers can be assembled and ranked.

The PCST problem is NP-hard, as a generalization of the basic Steiner Tree problem, but effective procedures for it are available (Ljubić *et al.*, 2006),(Bailly-Bechet *et al.*, 2011). Variants of the problem were previously used in bioinformatics, notably by E. Fraenkel’s group: Huang and Fraenkel (Huang and Fraenkel, 2009) applied a PCST model to transcriptomic, phosphoproteomic and genetic screen data to detect changes in regulatory and signaling pathways in yeast. Bailly-Bechet et al. used PCST to analyze transcription data related to pheromone response in yeast. Tuncbag et al. (Tuncbag *et al.*, 2013) used a prize-collecting Steiner forest (PCSF) formulation to discover multiple altered pathways induced by pheromone response in yeast using transcriptomic and proteomic data. Gitter et al. (Gitter *et al.*, 2014) generalized the PCSF problem to find a “consensus network” shared across multiple patients. Here we used an algorithm due to Akhmedov et al. (Akhmedov, LeNail, *et al.*, 2017) implemented in R (Akhmedov, Kedaigle, *et al.*, 2017) in order to solve PCST problems.

In testing on five TCGA cancer cohorts spanning >2500 patients and comparison to validated driver genes, PRODIGY outperformed extant methods and ranking based on network centrality measures. Our results emphasize the pleiotropic effect of driver genes and show that PRODIGY is capable of identifying even very rare drivers. Hence, PRODIGY can assist oncologists in decisions regarding personalized treatment.

### Caveats

Note that while we occasionally talk about driver mutations, all our analysis is done on the gene level and - as in SCS and DawnRank - different mutations in the same gene are not distinguished. Since the number of mutations per mutated gene in a patient is usually 1 (**STable 1**) this distinction is less important for personalized ranking than for cohort-level analyses. Also, as we shall see, often we identify and rank ten genes or more per patient, so the notion of drivers in this study is somewhat more lenient than is common in the literature. However, our results suggest that a larger number of predicted drivers actually contribute to the performance.

## Methods

Given the set of mutated genes and the expression profile of an individual, we wish to rank the mutated genes in that individual. Our assumption is that the influence of driver genes is disseminated along pathways and is manifested by DEGs. By aggregating evidence from multiple pathways for a mutated gene, we score the extent to which it explains the deregulation of the pathways. This score serves as a proxy to the likelihood that the gene is a driver in the patient. Mathematically, we score the influence of a mutation on a deregulated pathway using the undirected prize collecting Steiner tree (PCST) model.

### The PCST model

In this problem (**Figure 1A**) the goal is to find in a weighted graph a subtree maximizing the sum of the weights of the nodes minus the cost of edges in it. The input is an undirected graph *G* = (*V, E, W, P*). *W*: *E* → *R*_+_ is a positive weight function on the edges and *P*: *V* → *R* is a weight function on the nodes. In our context, edge weights are penalties reflecting interaction reliability, positive node weights are prizes given to DEGs, while other nodes that can serve as intermediate nodes in the tree (*Steiner nodes*) are assigned non-positive values serving as penalties. Given a node *g* ∈ *V*, the objective is to find a subtree *T* of *G* that contains *g* and maximizes:

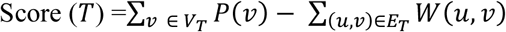

**Figure 1:**
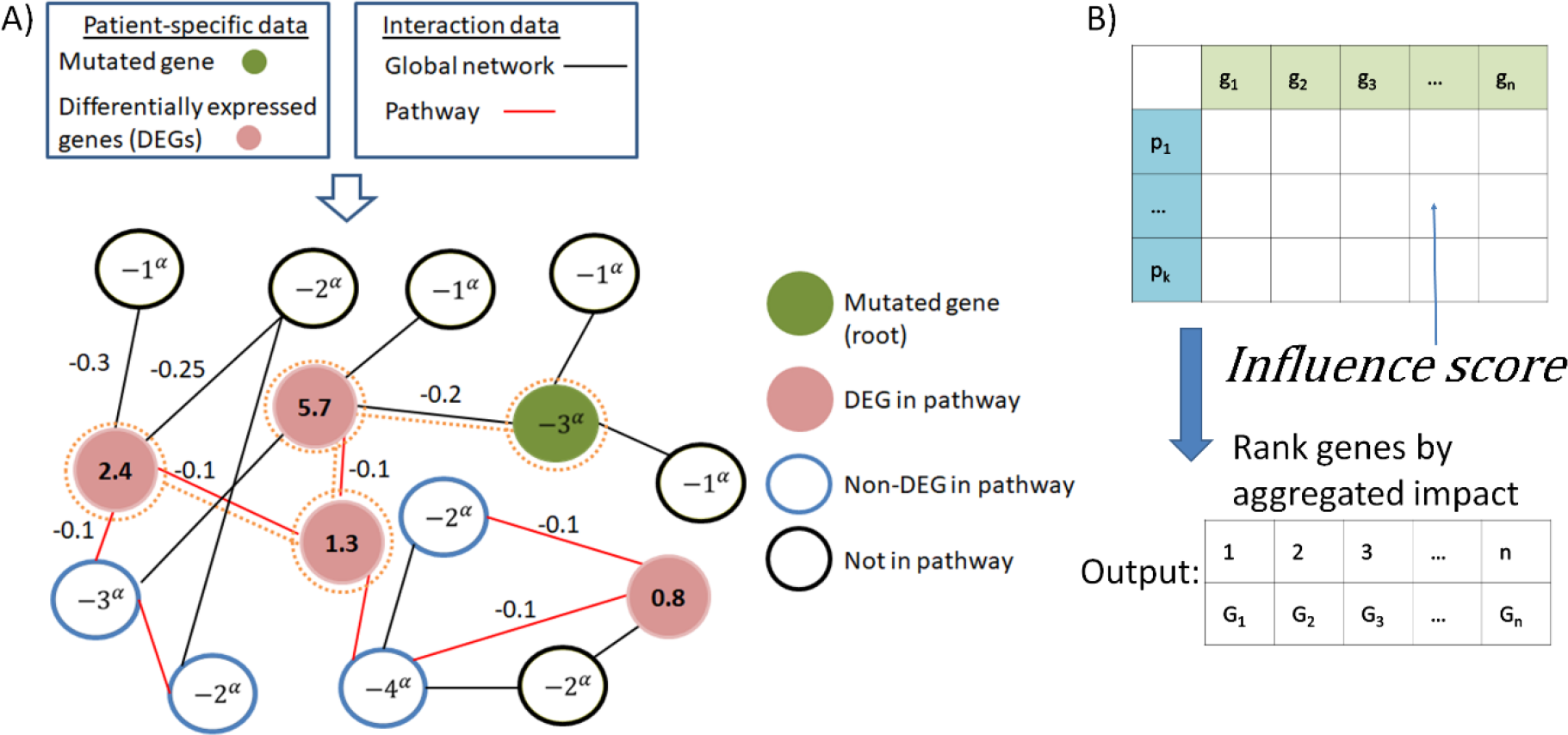
Outline of Prodigy’s approach. A) Scoring the influence of the mutated gene *g* on the pathway *p*: The pathway and genes at distance 1 from it or from the mutated gene *g* in the global network, along with the global edges among them, constitute the network *G*_*p,g*_ for analysis (see **Methods**). This is the network shown here. Node prizes (positive values) reflect the extent of differential expression of DEGs in *p*, and node penalties reflect other node’s degrees (calibrated by the exponent α). Edge penalties reflect interaction confidence. The goal is to find a maximum weight subtree in the network rooted at the mutated gene *g*. Its weight is the score of the PCST solution. In this example the subtree marked by orange dotted lines is the PCST solution, of score 9-3^α^. The *influence score* of the pair (*p,g*) is the score of the PCST solution, divided by the sum of the values of DEGs that belong to *p* (10.2 here). B) After calculating the influence score for all pairs (*p,g*), we filter out some pathways and genes from the scoring matrix (see **Methods**). The final output is a ranking of the remaining genes by their aggregated score on the remaining pathways.

In other words, the score of *T* is the total profit of pre-defined prizes minus the penalties of using intermediate edges and nodes. In our case, we also require that a specific mutated gene *g* is included in *T*. This way, *T* offers an explanation how the mutation in *g* causes the deregulation in the network: If *g* is a driver gene then a lot of the deregulation should be explainable by the tree.

### Data and reference network

Prodigy uses two types of genomic data for each patient: the list of mutated genes, i.e. all genes with SNVs or small insertions/deletions in coding regions, and the profile of mRNA expression. mRNA expression profiles from healthy tissue samples are also utilized for differential expression analysis. Prodigy also uses two types of undirected interaction networks: 1) a global PPI network taken from STRING v10.5 (Szklarczyk *et al.*, 2015). Here we used only physical interactions that were validated experimentally and interactions from other curated databases with confidence score > 0.7, so that only highly reliable interactions were included. The resulting network had 11,302 nodes and 273,210 edges. 2) A collection of pathways. Here we used either Reactome (Joshi-Tope *et al.*, 2005), NCI PID (Schaefer *et al.*, 2009) or KEGG (Ogata *et al.*, 1999). Information about the pathway databases is given in **STable 2**.

### The Prodigy algorithm

A schematic view of the algorithm is given in **Figure 1.** The algorithm works as follows:

#### Pre-processing

Given a patient’s mRNA expression profile (as read counts), differential expression analysis was done using DeSEQ2 (Love *et al.*, 2014) by comparing the profile to a background expression distributions from healthy samples of the same tissue of origin. All genes with > 2 log2-fold-change that are statistically significant (FDR = 0.05) were identified as DEGs.

The gene set of each pathway is tested for enrichment in DEGs using the hyper-geometric score, and pathways that are significantly enriched (FDR = 0.05) are called *deregulated*.

#### Driver - pathway scores

We use a global interaction network *G* = (*V, E, W*) where *W* is the edge confidence score. For a deregulated pathway *p* we also have its network *G*_*p*_ = (*V*_*p*_, *E*_*p*_). Both networks are undirected. The influence score of the mutated gene *g* on pathway *p* is calculated as follows:

1. We construct a new network *G*_*p,g*_= (*V*_*p,g*_, *E*_*p,g*_, *W*_*p,g*_, *P*_*p,g*_) that is derived from *G, G*_*p*_ and *g*, as follows: The nodes of the network are those of the deregulated pathway, *g*, and *N*(*V*_*p*_ ⋃ *g*)- their distance 1 neighbors in *G*:

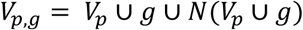 Its edges are those of the deregulated pathway plus all edges of the global network with both ends in *V*_*p,g*_:

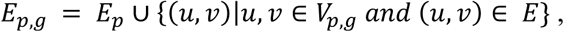 The cost of the edges from *p* is 0.1. For the other edges, which originate from the global network *G*, their cost depends on their confidence score in that network, with edges of higher confidence costing less.

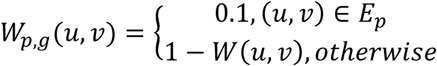 Edges from the pathway are assigned a constant penalty of 0.1 since pathway databases do not provide confidence scores for the interactions, but those pathways are highly curated. In contrast, the confidence scores on the edges from the global network are given an upper bound of 0.8 so that their cost in *G*_*p,g*_ is at least 0.2. The rationale is that we want to steer the algorithm to prefer the original pathway edges, while allowing some alterations. Finally every DEG that belongs to the pathway has a positive (prize) score depending on its fold change (FC), and every other node *ν* has a negative (penalty) score depending on its degree in *G*_*p,g*_ as follows:

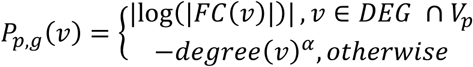 Note that DEGs in *V*_*p,g*_\*V*_*p*_have negative values. The PCST problem aims to collect as much of the prize nodes value while paying least penalty for intermediate edges and nodes. Intermediate nodes that have high degree (“hubs”) open more connection options and are thus penalized higher depending on their degree. The *α* parameter controls that penalty.
2. Having constructed *G*_*p,g*_ we now seek a tree *T*_*p,g*_ that contains *g* of optimal score. If Score(*T*_*p,g*_) ≤ 0 (i.e., no tree with positive score is found), an empty tree with score 0 is output instead.
3. To account for variability in pathway size and the number of DEGs in the pathway, the *influence score* of mutated gene*g* on pathway *p* is defined as the fraction of attained score out of the upper bound of all positive prizes in the pathway:

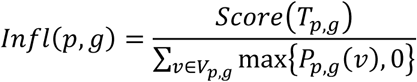

The overall *influence score* of *g* is ***infl***(***g***) = **Σ**_*p*∈*DP*_ *infl*(*p, g*) where *DP* is the set of deregulated pathways of the patient.

#### Pathway filtering

We compute driver-pathway influence scores for all mutated genes and all deregulated pathways. For the final score we exclude pathways for which more than half of the genes had a positive score. These are mainly very large pathways that have high connectivity in the global network, and therefore some genes may acquire positive influence scores by chance.

#### Gene filtering

Genes that acquired positive scores in many pathways have greater chance to represent a true effect on the tumor than genes that attained positive scores for only few pathways, possibly due to the topology of the network. In some patients, when plotting the distribution of *Infl*(*g*) scores across all mutated genes g (after filtering pathways), we observed a bimodal distribution (see **SFig. 2**). Typically, one distribution contains genes with high scores collected from many pathways and the other contains genes with low scores collected from a few pathways. We modeled this distribution as a mixture of two Gaussians and computed its maximum likelihood parameters using EM(Mc and Peel, 2000). We then excluded all genes that had higher posterior probability to come from the distribution with the lower mean (**SFig. 2**). If the fitted bimodal distribution had lower likelihood than a fitted unimodal normal distribution, we did not filter any gene.

#### Final ranking

After the filtering steps, genes are ranked according to their overall influence scores.

### Comparison to other personalized methods

We compared Prodigy to DawnRank (Hou and Ma, 2014) and SCS (Guo *et al.*, 2018). Both DawnRank and SCS used the global directed network taken from Wu et al (Wu *et al.*, 2010), with 11,648 nodes and 211,794 directed edges, so we tested Prodigy using as the global network an undirected version of the Wu et al. network, as well as using the SPRINT network. To gauge the effect of the topology of the network, we also generated personalized rankings using three node centrality measures: node degree, closeness and betweenness (see **Supp. Methods** for definitions). To produce rankings based on each measure, we calculated it on each of the networks *G*_*p,g*_ and summed the results over all the networks for a final ranking. Comparison with Hit’nDRIVE was not possible, as it outputs an unranked set of drivers for each individual and our performance measures rely on the ranking.

### Validation

In order to validate rankings, we used a curated list of driver genes from the Cancer Gene Census (CGC) as gold standard. CGC is part of COSMIC (Forbes *et al.*, 2017), the largest database of somatic mutations in cancer. CGC contains mutations of different forms (gene amplifications, SNVs, translocations etc.) that were experimentally validated as driver mutations for different cancer types. Since we only used information about SNVs and short indels of each patient, we used as ground truth only genes that were classified by CGC as containing a driver SNV or indel (n = 248 out of 567). In this validation, we assumed that if a gold-standard gene was mutated in a patient, it is a true driver gene in the patient’s tumor. We measured the quality of each method by means of precision, recall and F1 with respect to the gold standard (see **Supp. Methods**)

### Driver-Pathway linkage

Prodigy can quantify driver-pathway associations, allowing us to explore novel interactions and even cancer subtype-specific ones. Our hypothesis was that if driver gene *g* often deregulates pathway *p* then they will be observed together more frequently in patients of the cohort, and the deregulation state of *p* will be higher when *g* is acting as a driver. To test this conjecture, we focused on the ten top ranked genes for each individual and looked for driver-pathway pairs where the number of patients for whom the gene was ranked high and the pathway was deregulated was unexpectedly high according to the hyper-geometric distribution. For each pair, we then tested if *p* was more deregulated when *g* was classified as driver using t-test (see **Supp. Methods** and **SFig. 4** for more details).

## Results

### Driver gene ranking

We tested six ranking methods on 2569 samples from five cohorts of cancer patients from TCGA: COAD, LUAD, BRCA, HNSC and BLCA (Koboldt *et al.*, 2012; Weinstein *et al.*, 2014; Muzny *et al.*, 2012; Lawrence *et al.*, 2015; Collisson *et al.*, 2014) (212, 487, 969, 502 and 399 samples, respectively). Prodigy has a single tunable parameter - the node degree penalty *α* - and in order to choose its value, we used a training set comprised of 10% of the samples in each cohort to derive the *α* value giving highest F1 score. We then used that value (*α* = 0.05) to calculate personalized rankings for the remaining 90% for all cohorts and all pathway sources. The personalized rankings are available from the authors upon request. Prodigy’s results were stable across a range of *α* values with notable decline for *α* > 0.2 (**SFig. 4**).

**Figure 2A** shows the average precision, recall and F1 for Prodigy, DawnRank and for the three centrality measures using the Reactome pathways (see **Methods**). The results are reported as average values for the entire cohort as a function of the top N ranked genes. If an individual had less than N ranked genes, the last value for this patient was duplicated so that all quality measure vectors for all patients are of length N. Since SCS reported empty rankings for 720 samples (28%), it is not shown in **Figure 2A**. Performance of all methods on the set of patients for whom SCS produced results (the “SCS sub-cohort”) is shown in **SFig. 6**, and performance for different cancer types is shown in **SFig.7-9**. Results for the KEGG and NCI pathway databases for the entire cohort were similar (**SFig. 5**).

**Figure 2:**
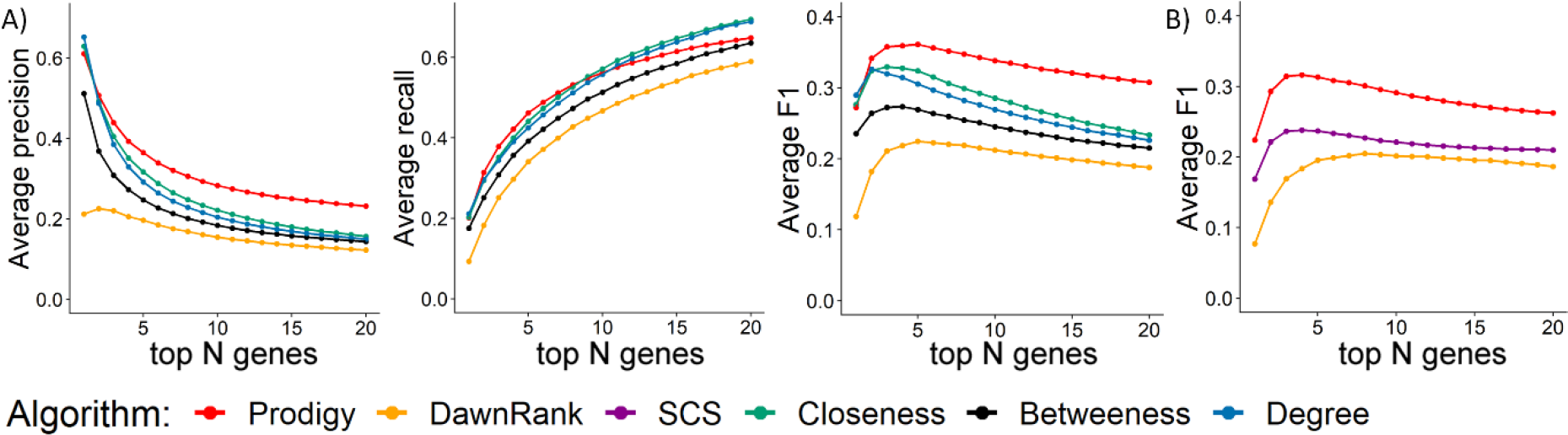
A. Average precision, recall and F1 across all patients (n=2340) as a function of the number of top ranked genes in the personalized profiles. Prodigy’s results were derived using the STRING global PPI network (see **Methods**) and Reactome pathways B. Average F1 using the global network from the SCS and DawnRank papers, on those patients for whom SCS proposed drivers (n=1804).

Overall, Prodigy outperformed SCS and DawnRank in terms of F1, precision and recall. SCS was better for the SCS sub-cohort on the NCI pathways from the ninth gene up and for high values of N on KEGG pathways. More generally, Prodigy showed improvement in performance as the number of tested pathways grew: NCI had fewest pathways, KEGG had more and Reactome most, and Prodigy’s results were weakest for NCI and strongest for Reactome. We explore this point further below.

To ensure that the improvement in results does not stem from the difference in the underlying networks, we also tested Prodigy on the same network used by DawnRank and SCS with two adjustments: (1) Since Prodigy works on undirected graphs, we ignored edge directions. (2) Since this network is unweighted, we gave weight=0.2 to all edges (and 0.1 to pathway edges as before, see **Methods**). The results (**Figure 2B** and **SFig. 10**) clearly show that Prodigy outperforms DawnRank and SCS even on their network.

Remarkably, the centrality measures produced very good predictions, consistently better than DawnRank and SCS – but worse than Prodigy. These measures had better recall than Prodigy, probably due to the fact that no filtering was done on the centrality measures while Prodigy excluded genes not likely to be drivers for an individual. The fact that driver genes are associated with high network connectivity was previously discussed (Shrestha *et al.*, 2017),(Jonsson and Bates, 2006),(Porta-Pardo *et al.*, 2015) and we observed it as well: in our global network derived from STRING, known drivers included in the CGC tended to have high degree and betweenness (**SFig. 11**). Our results emphasize the need to account for “hubness” property in methods for driver gene ranking. Prodigy accounts for this factor by penalizing Steiner nodes according to their degree. Taken together the results clearly demonstrate that Prodigy outperforms mere topology measures in capturing true driver genes.

### Comparison to a cohort-level method

We wished to compare Prodigy to a state-of-the-art method for cohort-based identification of driver genes. For this purpose, we chose MutsigCV (Lawrence *et al.*, 2013), a leading algorithm for that purpose, and performed the comparison on a cohort of 178 LUSC patients from the TCGA. MutsigCV produced cohort-level ranking of drivers *ρ* on that data. To produce a ranking based on *ρ* for each patient, we computed the ranking of the individual’s mutated genes induced by *ρ*. Mutated genes not included in *ρ* were excluded. Prodigy was applied to each patient using Reactome pathways and *α* = 0.05. The results (**SFig. 12**) show that Prodigy outperformed MutsigCV in terms of precision, recall and F1 from the gene ranked fourth up. Importantly, Prodigy identified rare drivers that ranked very low by MutsigCV (**STable 3**). Overall, 37 known drivers from the CGC were mutated in less than 2% of the LUSC cohort, identified by Prodigy as personalized drivers (rank < 10), and ranked > 500 among the cohort-level genes according to MutsigCV. This illustrates Prodigy’s ability to pinpoint even very rare drivers, which are missed by cohort level methods.

### Distinguishing true mutations from decoys

In order to test Prodigy’s ability to distinguish drivers from non-drivers, we examined its discrimination power between top ranked drivers and a random set of decoy mutations (see **Supp. Methods**). The results (**SFig 13**) show that Prodigy can discriminate quite well drivers from decoys.

### Discovering rare drivers

One of the advantages of Prodigy is its ability to identify rare drivers, even when the gene is mutated in few patients, as shown in the comparison to MutsigCV. To further demonstrate this ability we looked for genes that were mutated in < 2% of the respective cohort and were ranked in the top 10 drivers of individuals. The results are summarized in **Figure 3**. In some cohorts, most of the mutated genes were in fact rare (< 2%, **STable 4**), which is of course reflected in our results. On the other hand, Prodigy prioritized rare mutations to lesser extent than their frequency in the population (**STable 4**). Moreover, Prodigy was capable of identifying known drivers from the CGC that were rarely mutated in the cohort (**STable 5**). Taken together, this demonstrates Prodigy’s ability to detect both rare and frequent drivers.

**Figure 3:**
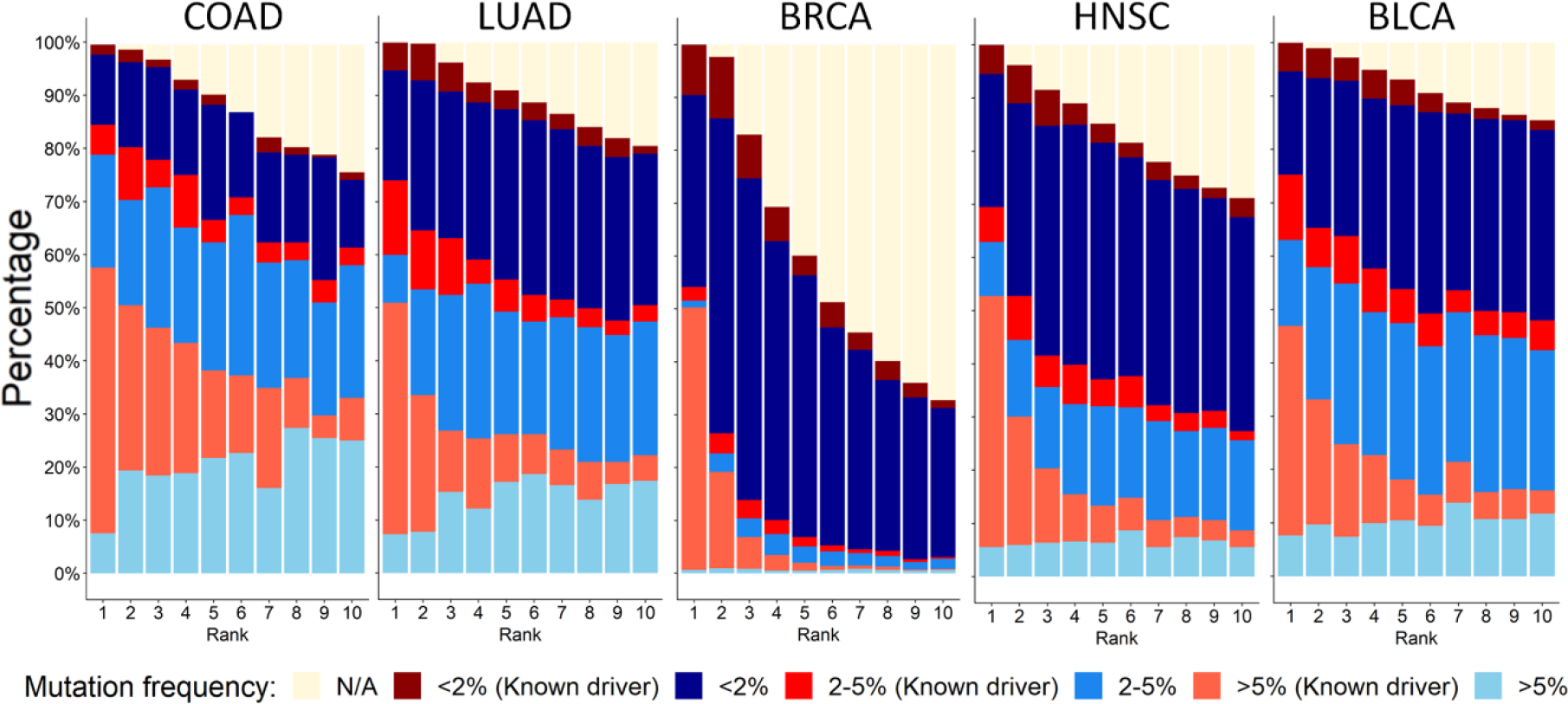
Prodigy discovers rare drivers. For each cancer type and for each individual we analyzed the top 10 genes according to the ranking. For k = 1,…,10 the plot shows the fraction of patients for whom the gene ranked k-th belongs to the respective frequency bin (as denoted by its color). N/A: patients for whom Prodigy ranked less than k mutations.

### Driver gene-pathway linkage

We identified 1299 significant driver-pathway interactions (see Supp. File 1). They include some very well-known interactions between *TP53* and sub pathways of the cell cycle in all cohorts except COAD and *TP53*-DNA repair pathways in the BLCA cohort. Moreover, the gene *A2M* was associated with “G alpha (i) signaling events” in the COAD, BRCA and BLCA cohorts. The G alpha (i) signaling pathway belongs to the GPCR family of signaling pathways, which are strongly linked to cancer (Dorsam and Gutkind, 2007). This analysis can provide new insights on the mechanism by which the drivers operate and can offer new targets for further research.

### Multi-pathway effect

One of the main assumptions underlying Prodigy is that driver genes affect cellular process pathways, and therefore summarized scores from multiple pathways will improve our ability to identify them. This is in contrast to previous methods that took a global approach to driver gene prioritization based on a single unified picture of the state of the tumor^22,23^. In order to test whether multiple sources indeed contribute to the accuracy of prediction, we explored the performance as a function of the number k of allowed pathways per mutated gene. For k = 1,…,50, we used the top k scoring pathways of each gene for ranking and examined the average area under the precision-recall curve (AUPR) for each cohort (see **Supp. Methods**). **Figure 4A** shows that for all cohorts, AUPR improved with incorporating more pathways and plateaued at 15-30 pathways.

**Figure 4:**
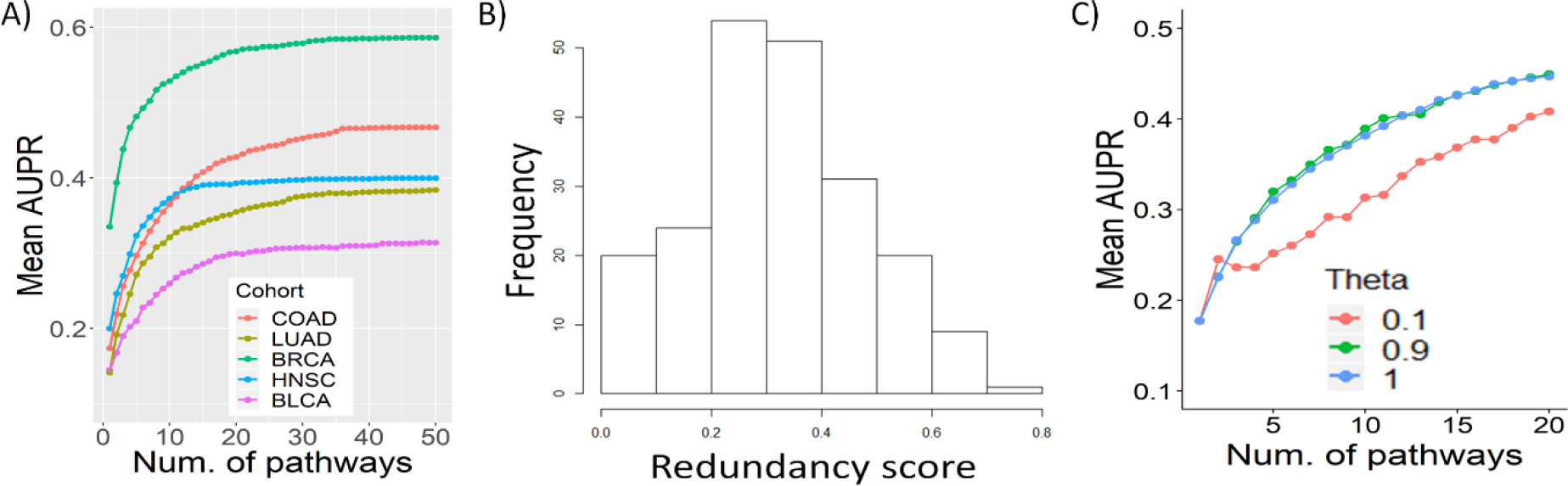
Multi-pathway effect: A) Mean AUPR as a function of the number of top scoring pathways per gene used to derive the results. B) The distribution of redundancy between the top 20 pathways per patient in the COAD cohort (n = 212, see **Supp. Methods**). C) Removal of pathway redundancy. The plot shows the AUPR for predicting driver genes in the COAD cohort when filtering out overlapping pathways among the top scoring pathways per gene (**Supp. Methods**). *θ* is the maximum allowed Jaccard Index between included pathways (*θ* = 1 implies no filtering).

Since different pathways may partially overlap, we tested the extent of this overlap and its effect on performance. We computed the distribution of Jaccard Index between pairs in the top 20 scoring pathways of each gene (i.e., the number of genes that belong to both pathways divided by the number of genes in their union, **Supp. Methods**). The results show substantial overlap among the pathways that contribute to the rankings (**Figure 4B**). However, when we filtered out such overlapping pathways, assuming they contain the same information and thus unnecessary for accurate prediction, performance only moderately degraded (**Supp. Methods** and **Figure 4C**). Taken together, we demonstrated the usefulness of using multiple pathways in order to rank driver genes, even when there are overlaps among them.

### Actionable and druggable targets

Prodigy’s rankings can aid the oncologist in deciding on a patient’s therapy, by matching treatment to the predicted driver genes. In order to explore this possibility we used two data sources: (1) DGIdb 3.0 (Cotto *et al.*, 2017), which contains drug targets (or *druggable genes*, i.e., genes with directed pharmacotherapy). Here we used only cancer-specific sources from DGIdb and identified 1375 genes. (2) TARGET (Van Allen *et al.*, 2014), which lists *actionable genes* (i.e., genes for which a genomic-driven therapy exists). The total number of actionable genes was 60. We explored not only the ranked mutated genes themselves but also the pathways that were highly linked (influence score > 0.8) to at least one gene of the top 10 ranked genes of an individual. The rationale is that these pathways are most altered by the driver genes and thus can be targeted in potential treatments. The results (**Figure 5)** indicate that most patients harbor at least one druggable driver (a druggable gene that was prioritized as a driver by Prodigy; mean: 3.32, sd: 2.01) but many do not contain any actionable drivers (mean: 0.82, sd: 0.87). As expected, the number of target genes increased substantially when genes from highly linked pathways were also considered. More importantly, the number of patients without any druggable or actionable gene decreased below 10%. The only exception was the HNSC cohort, where the number of patients without actionable genes remained high (35.8%) even when considering pathways. Hence, Prodigy is able to suggest possible therapeutic targets personally tailored to the patient’s driver genes and uses information about the pathways that are deemed altered by the drivers in order to expand pharmacotherapy options.

**Figure 5:**
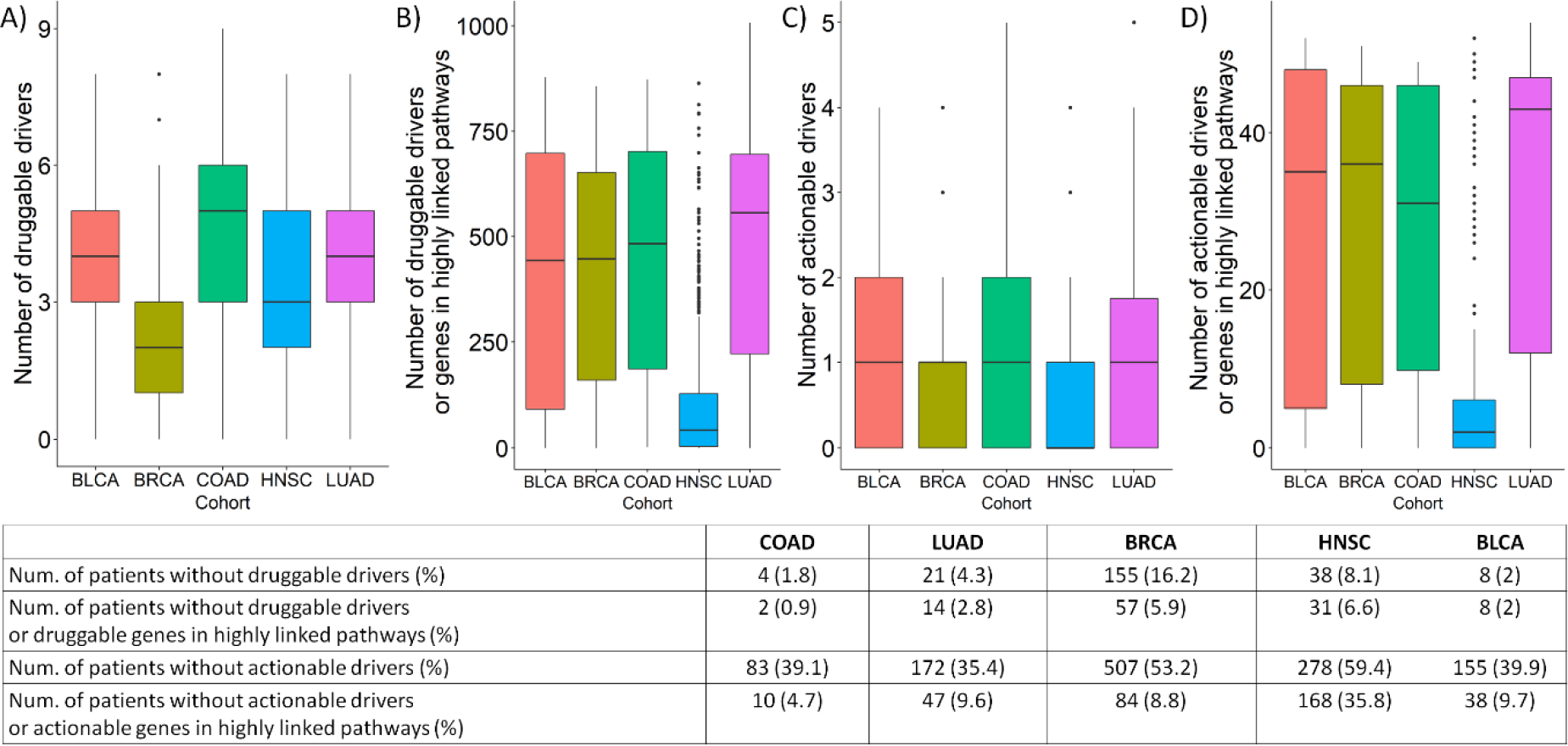
Actionable and druggable genes. The box plots show the distribution of the number of actionable and druggable genes (i.e. genes from TARGET^43^ and DGIdb^42^) per patient across the different cohorts. A and C: The distribution of the number of druggable and actionable drivers among the 10 top genes ranked by Prodigy. B and D: The distribution of the number of druggable and actionable genes among predicted drivers and their highly linked pathways. The table describes the number of patients without any druggable/actionable genes in the four categories with respect to the cohort.

### Implementation

Prodigy was implemented in R and the software is available in https://github.com/Shamir-Lab/PRODIGY. Mean runtime was about 5 minutes per patient on a 65 core, Intel(R) Xeon(R) 2.30GHz, 755GB RAM server.

## Discussion

Personalized diagnosis of cancer patients is a precondition for planning treatment. Deciphering the altered mechanisms and the mutated genes driving them gives a comprehensive picture of the state of the tumor. Although many driver mutations were experimentally validated, there is great potential benefit in identifying which genes act as drivers *in an individual* and prioritizing them: Driver genes are diverse even within a cancer subtype, and they may be rare or not match the disease organ.

Here we provide a novel algorithm called Prodigy for driver gene prioritization in an individual based on the patient’s tumor mutation and expression profiles. In testing of over 2500 patients from five cancer subtypes, Prodigy substantially outperformed prior methods for the task. All methods (including ours) use an underlying global interactions network, and we observed that using simple centrality measures of that network to prioritize genes gives better results than the prior methods. This is probably since driver genes tend to have higher connectivity in large PPI networks (**Supp. Fig 11**). Prodigy is able to overcome this bias, as manifested by its advantage over centrality-based prioritization. This is achieved by penalizing for high degree of Steiner nodes. It is important to emphasize that Prodigy does not utilize any information regarding the tissue from which the tumor originated, hence the reported summary of results by cancer type is somewhat arbitrary, and the aggregated results are more meaningful. A limitation of our analysis and of prior studies was the use of the CGC collection of (global) driver genes for validation. It served in lieu of a gold standard, since no experimental patient-specific driver information was available.

A unique feature of Prodigy is the fact that it collects information from many distinct pathways in the analysis of a patient. Our results shed light on the pleiotropic effect of drivers on individual tumors, since we showed that highly ranked driver genes tend to exert influence across many pathways. This hypothesis was demonstrated and discussed before (Vogelstein *et al.*, 2013; DeSimone *et al.*, 2013; Wang *et al.*, 2016), and Prodigy might help in systematic analysis of this phenomenon. However, such application requires further biological validation that is beyond the scope of this work. Furthermore, we demonstrated that this pathways perspective is more powerful than previous approaches that utilized one global network for the analysis. While Prodigy ranks the genes without setting a cutoff for driver detection, our analysis shows the F1 scores peak around N=5. On the other hand, recall rises for N>5, so by using prior knowledge about driver genes and observing the actual influence scores of genes that are ranked lower, additional drivers can be pinpointed and used.

Our analysis shows that Prodigy can identify even very rare driver genes, and can reveal linkage between a driver gene and pathways that are preferentially deregulated when the gene acts as a driver. The identified genes typically have multiple drug targets, and thus can suggest treatments. A limitation of Prodigy is its relatively high running time. While this is not a concern when analyzing a single patient with a limited number of mutations, it prohibits analyzing thousands of mutations and obtaining empirical p-values by permutation tests.

## Acknowledgements

We thank Nimrod Rappaport for helpful comments on the manuscript. The results published here are based upon data generated by The Cancer Genome Atlas managed by the NCI and NHGRI. Information about TCGA can be found at http://cancergenome.nih.gov. Study supported in part by the Israel Science Foundation (grant No. 1339/18), by the DIP German-Israeli Project cooperation, and by Len Blavatnik and the Blavatnik Family foundation. GD was supported by a fellowship from the Edmond J. Safra Center for Bioinformatics at Tel Aviv University.

