## Supplementary Material for "PRODIGY: personalized prioritization of driver genes"

**Supplementary methods**:

Centrality measures:

Closeness $(v) = \frac{1}{\sum_{u \neq v} d(v,u)}$ where $d(v,u)$ is the weighted length of the shortest path from $v$ to $u$ in the graph.

Betweenness $(v) = \frac{\sum_{i \neq v \neq j} \sigma(i,v,j)}{\sum_{i \neq j} \sigma(i,j)}$ where $\sigma(i,j)$ is the number of weighted shortest paths between $i$ and $j$ and $\sigma(i,v,j)$ is the number of weighted shortest paths between $i$ and $j$ that pass through $v$.

Precision, recall, F1: Let $rank_i[k]$ be the set of top k ranking genes for the patient $i$, then

$$\text{Precision } (rank_i[k]) = \frac{|rank_i[k] \cap CGC\_genes|}{k}$$

Recall $(rank_i[k]) = \frac{|rank_i[k] \cap CGC\_genes|}{|SNV_i \cap CGC\_genes|}$ where $SNV_i$ is the set of mutated genes in patient $i$

$$\text{F1 } (rank_i[k]) = 2 * \frac{\text{Precision } (rank_i[k]) * \text{Recall } (rank_i[k])}{\text{Precision } (rank_i[k]) + \text{Recall } (rank_i[k])}$$

In order to calculate the quality of prediction for a cohort of patients, we averaged these quality measures over the entire cohort and showed the results as a function of $k$.

Driver-Pathway linkage identification pipeline:

1) First, we extract all driver-deregulated pathway pairs $(g, pw)$ that are observed together in more than three patients. Here we defined the set of drivers for each patient as the 10 top ranked genes.

2) For each driver-pathway pair $(g, pw)$ let $n_g$ be the number of patients for whom the gene $g$ was predicted as driver by Prodigy, $n_{pw}$ the number of patients for whom the pathway $pw$ was deregulated and $n_{g,pw}$ the number of patients for whom $g$ was predicted as a driver and $pw$ was deregulated. Let $N$ be the number of patients in the cohort. The probability of observing $n_{g,pw}$ patients is given by $hypergeometric(N, n_g, n_{pw}, n_{g,pw})$. We calculate the p-value for $n_{g,pw}$ of each pair and identify the *significant pairs* (FDR < 0.05).

3) For each significant pair $(g_i, pw_i)$, we perform a comparison between the absolute deregulation of $pw_i$ in patients for whom $g_i$ was predicted to be a driver and in patients for whom $g_i$ was not predicted to be a dirver using t-test (**SFig 4**). All pairs with statistically significant difference (FDR < 0.05) were classified as *significant interactions.*

4) To ensure that these interactions could not have been identified by observing the mutational state of the gene alone (i.e., regardless of its ranking), we ran the entire process on all mutated genes $g$ to identify *mutation-pathway* interactions. We then excluded significant driver-pathway interactions that overlapped significant mutation-pathway interactions. The driver-pathway pairs that passed this filter are reported as output.

Multiple-pathways effect: We repeated the following procedure for each value of *k* between 1 and 50. For each gene $g$ we used only the *k* pathways with the highest scores to calculate the gene's influence score $infl(g)$, and ranked all genes according to the score. We then computed the AUPR for the ranked list.

In order to examine the extent of overlap between the top *k* scoring pathways for each gene $g_i$, we computed for every pathway $pw_i$ in the top *k* pathways its maximum overlap with any other pathway among the top *k*, and averaged these scores over all the pathways as follows:

$$Overlap\ Index(g_i) = \frac{1}{k}\sum_{i=1}^{k} \max_{pw_i \neq pw_j} JI(pw_i, pw_j).$$

Here JI indicates the Jaccard Index, namely the number of genes shared by the two pathways divided by the number of genes in their union. We then averaged these values over all drivers to get the *Redundancy score*, a value that represents the extent of pathway overlap for the patient.

To explore the effect of this redundancy on the performance of Prodigy, we built the set of top pathways from which we derive the ranking in a sequential manner while avoiding redundancy: for each gene $g$ we ranked the pathways by their scores $infl(p, g)$ in decreasing order and added the next pathway to the set only if it had JI < $\theta$ with every pathway already in the set. We did so for all the genes mutated in the patient, ranked the genes according to their final influence score and calculated AUPR for the patient. The results in **Figure 5C** show the mean AUPR across all patients as a function of $\theta$ and *k.*

**Distinguishing true mutations from decoys**:

In order to further test Prodigy's ability to distinguish drivers from non-drivers, we examined its ability to discriminate the top ranked drivers from a random set of decoy mutations: Given a sample that contained $n$ mutated genes, we randomly selected $n$ other (non-mutated) *decoy genes*. Decoy gene $i$ was selected such that its degree was equal or higher than that of the truly mutated gene $i$, in order to control for possible degree bias in true drivers. We then ran Prodigy on the set of decoy mutations and examined the discrimination between true and decoy mutations according to their total influence scores in terms of ROC AUC. We ran this process for all 212 COAD patients from the TCGA using Reactome pathways and setting $\alpha = 0.05$.

We used the influence scores to rank the 10 top true mutations together with the k top decoy mutations for k=10,20,…,$n$, and computed the median AUC ROC for each value of k. The results are shown in **SFig 13**.

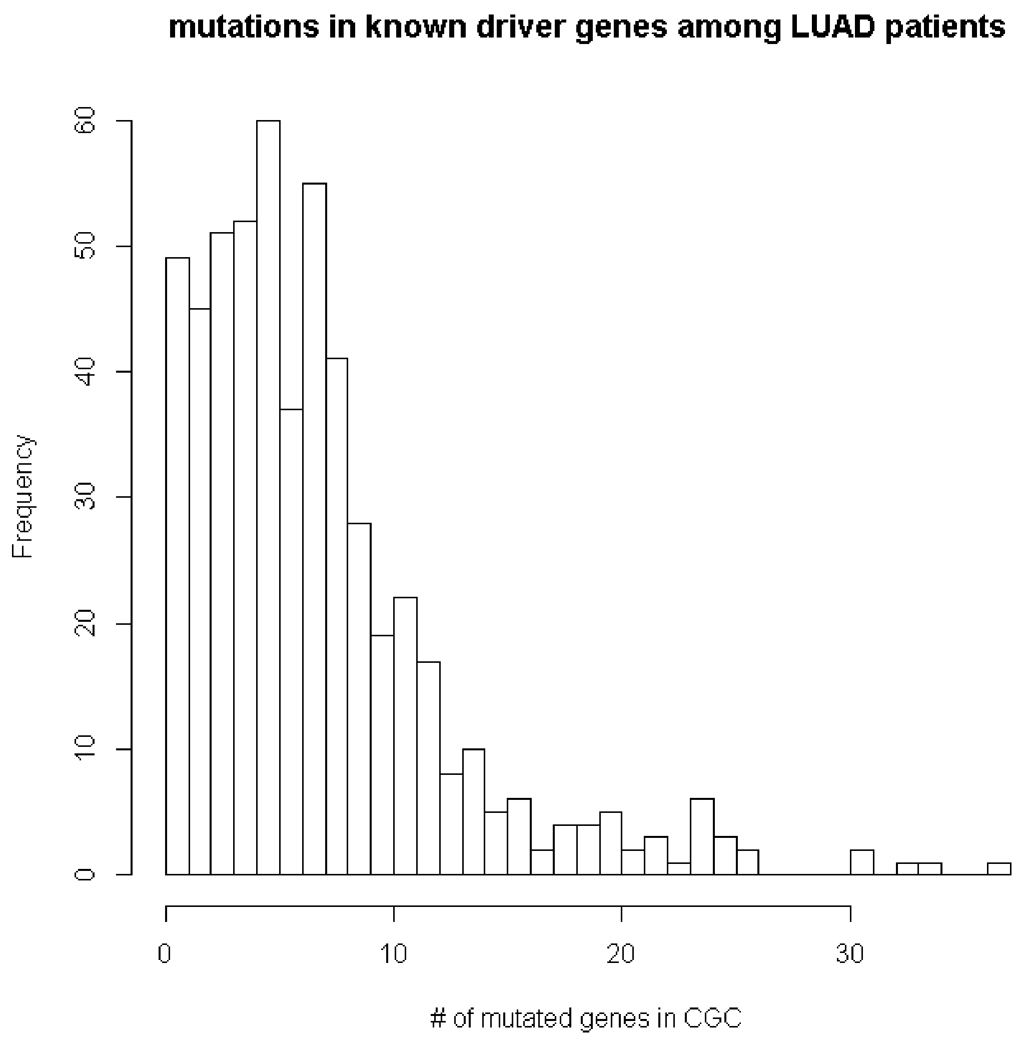


**Supplementary Figure 1**: Distribution of the number of mutated genes that are also in CGC[3] per patient, in the LUAD cohort.

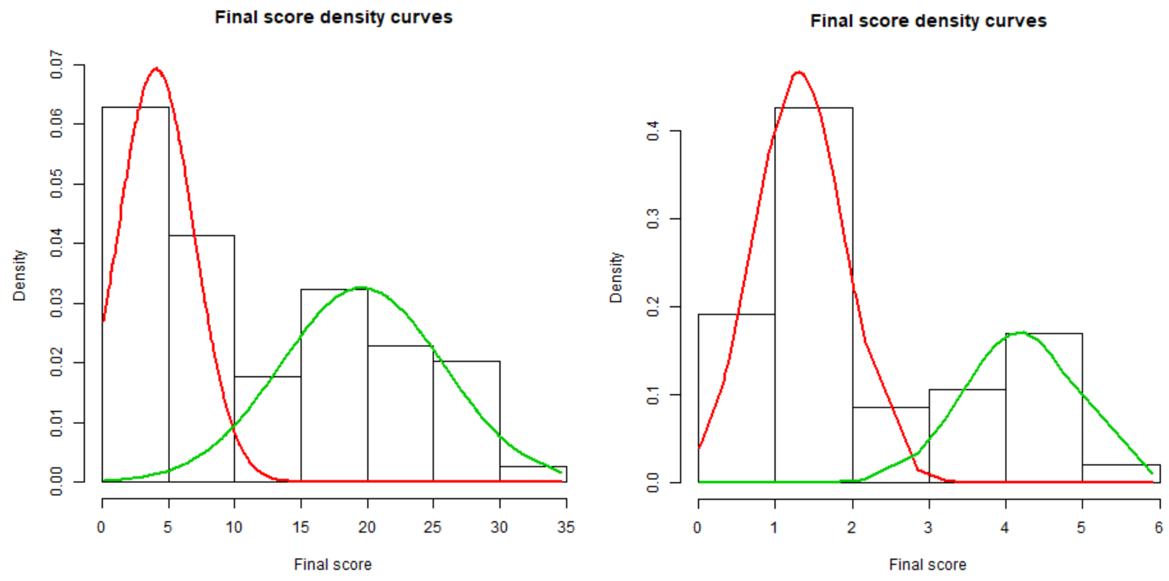


**Supplementary Figure 2**: Distribution of gene influence scores. The color lines show breakdown of the distribution into two Gaussians as detected by the EM algorithm. The red distribution reflects mutations that gain sporadic or low scores for few pathways due to the topology of the network. The green distribution reflects mutations with consistent scores across many pathways.

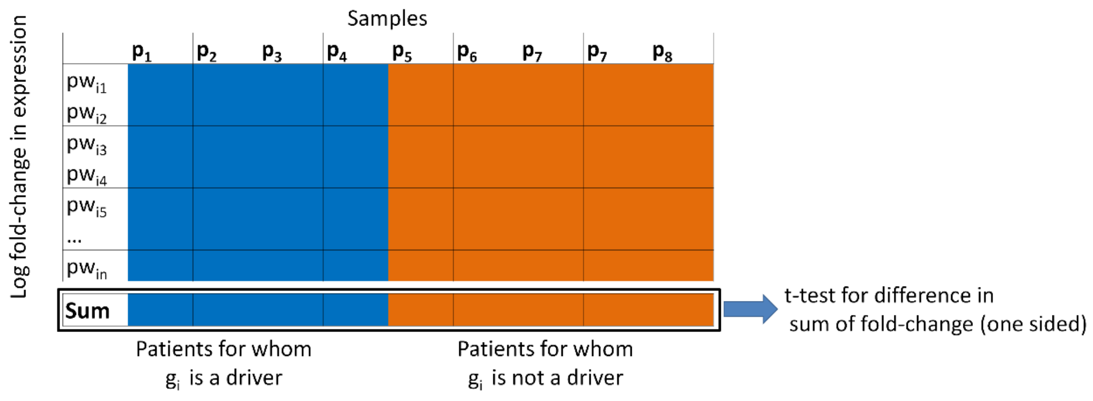


**Supplementary Figure 3**: Let $(g_i, pw_i)$ be a significant pair. The matrix contains the log2 fold-change in the expression of genes that belong to $pw_i$ (the genes $\{pw_{i1}, \dots, pw_{in}\}$). Columns in the blue / orange part of the matrix represent patients for whom $g_i$ was / was not identified as a driver by Prodigy. A t-test is performed based on the total sum of fold-changes for each patient. The null hypothesis is that the mean sum of fold-changes is not greater in the group of patients for whom $g_i$ was classified as a driver.

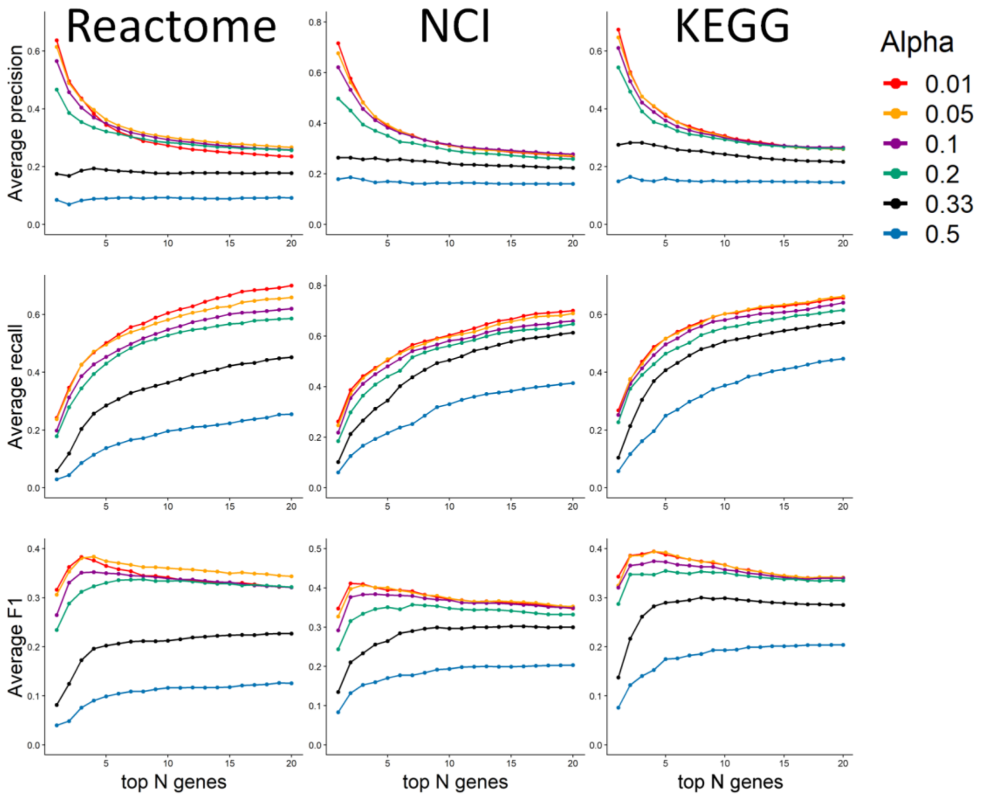


**Supplementary Figure 4:** Effect of the parameter $\alpha$ on Prodigy's results. The plots show average precision, recall and F1 on the training cohort (n=215) as a function of $\alpha$, the exponent of the penalty term for Steiner nodes and N, the number of top ranked genes. The training cohort was comprised of 10% from each of the five datasets used.

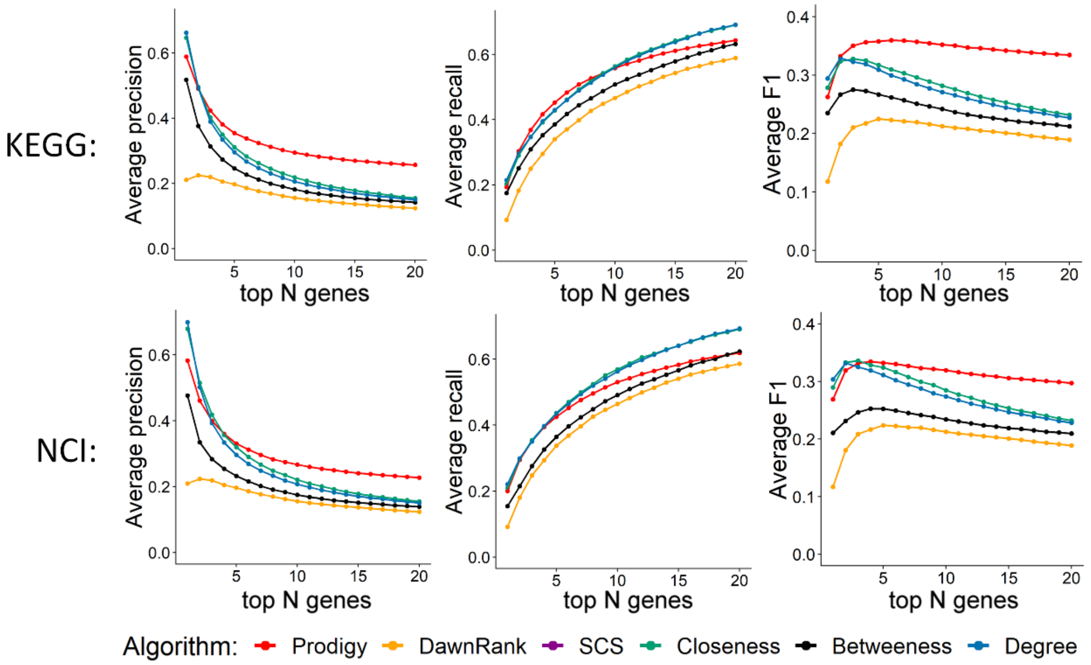


**Supplementary Figure 5**: Average precision, recall and F1 across all patients as a function of the top N genes in the personalized profiles. Results are for KEGG and NCI as pathway databases. Prodigy produced empty rankings for 14, 6, and 6 samples (<0.5%) in Reactome,

KEGG and NCI, respectfully, due to lack of deregulated pathways or zero influence scores for all genes.

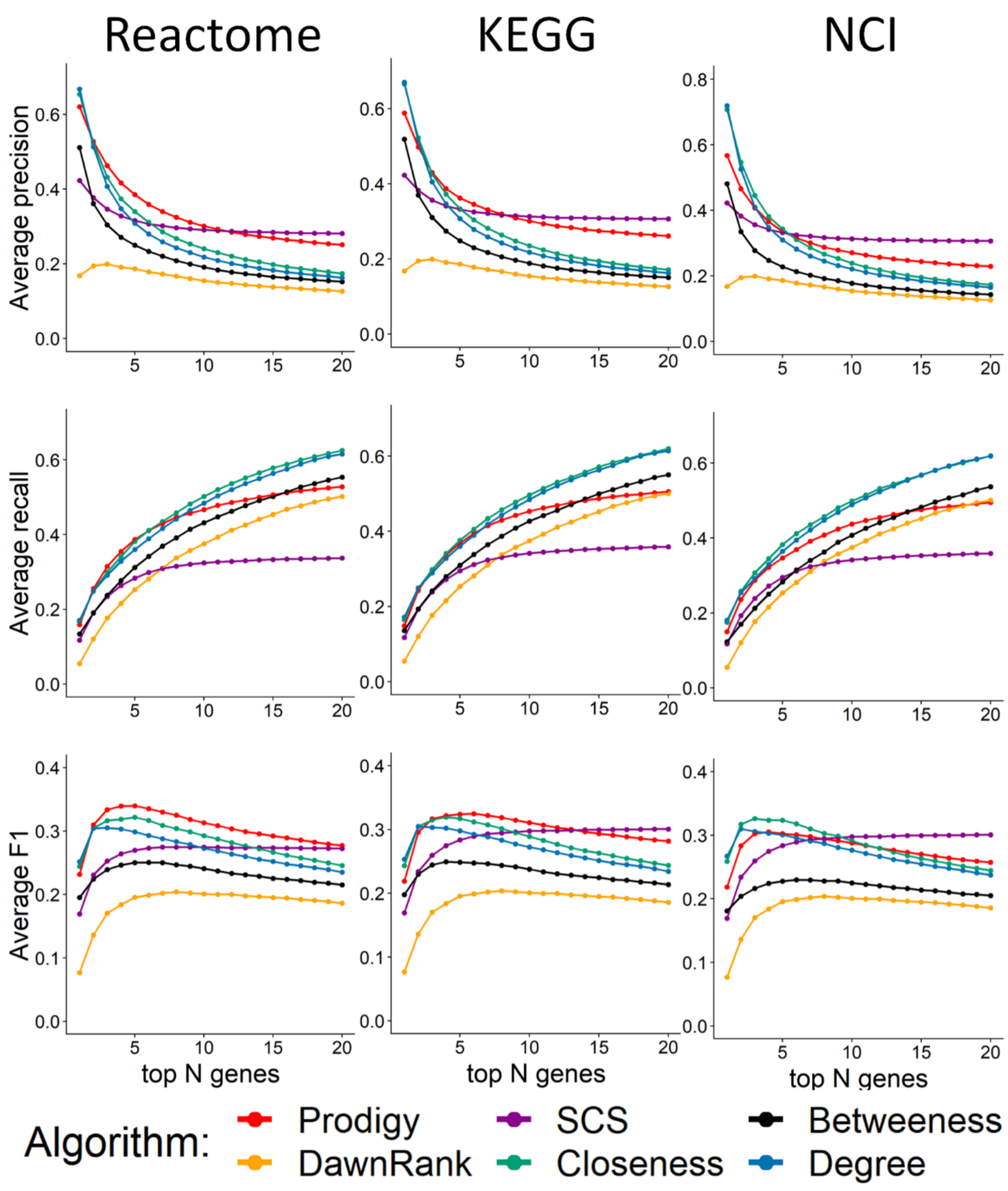


**Supplementary Figure 6**: Average precision, recall and F1 across all cohorts are presented as a function of the top N genes in the personalized profiles. The results are for the subset of samples for which SCS produced non-empty profiles (n=1847, 1849, 1849 for Reactome, KEGG and NCI).

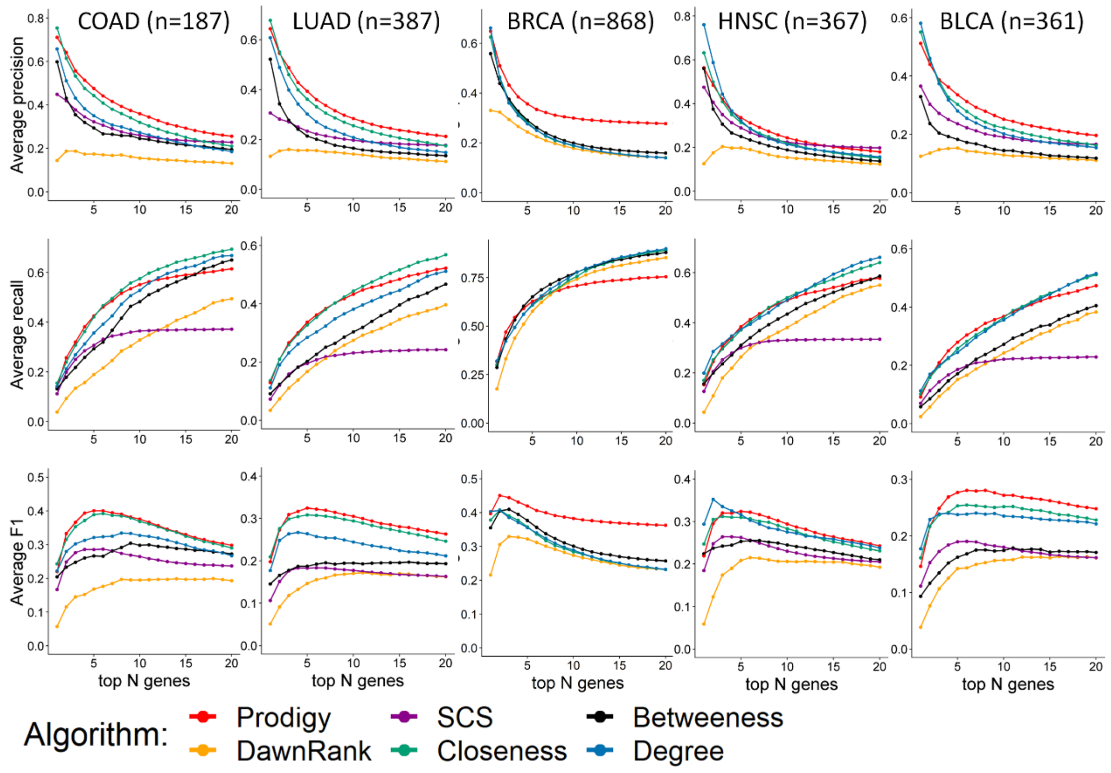


**Supplementary Figure 7**: Performance of the methods on each cohort using Reactome pathways. Results are shown for Prodigy, DawnRank, SCS and three centrality measures. The results of Prodigy and of the centrality measures were derived using STRING as global network and Reactome as pathway DB. SCS and DawnRank used their directed network. Average precision, recall and F1 across the relevant cohort are presented as a function of the top N genes. Results are for the "SCS sub cohort" (see **Methods**), except for the BRCA cohort where SCS produced empty rankings for 529 (55%) patients, hence it was excluded from the comparison. $\alpha = 0.05$ was used for all cohorts. n = sample size of the test group.

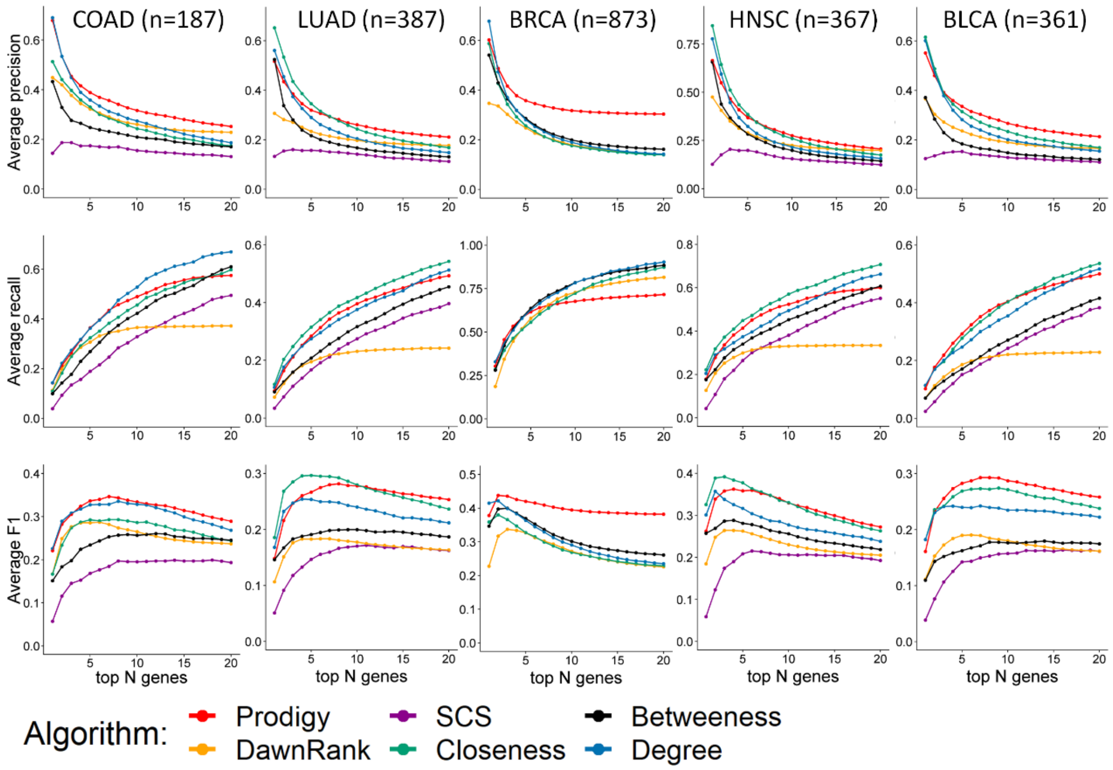


**Supplementary Figure 8**: Performance of the methods on each cohort using KEGG pathways. Details are as in Supplementary Figure 7 but here the KEGG pathway DB was used.

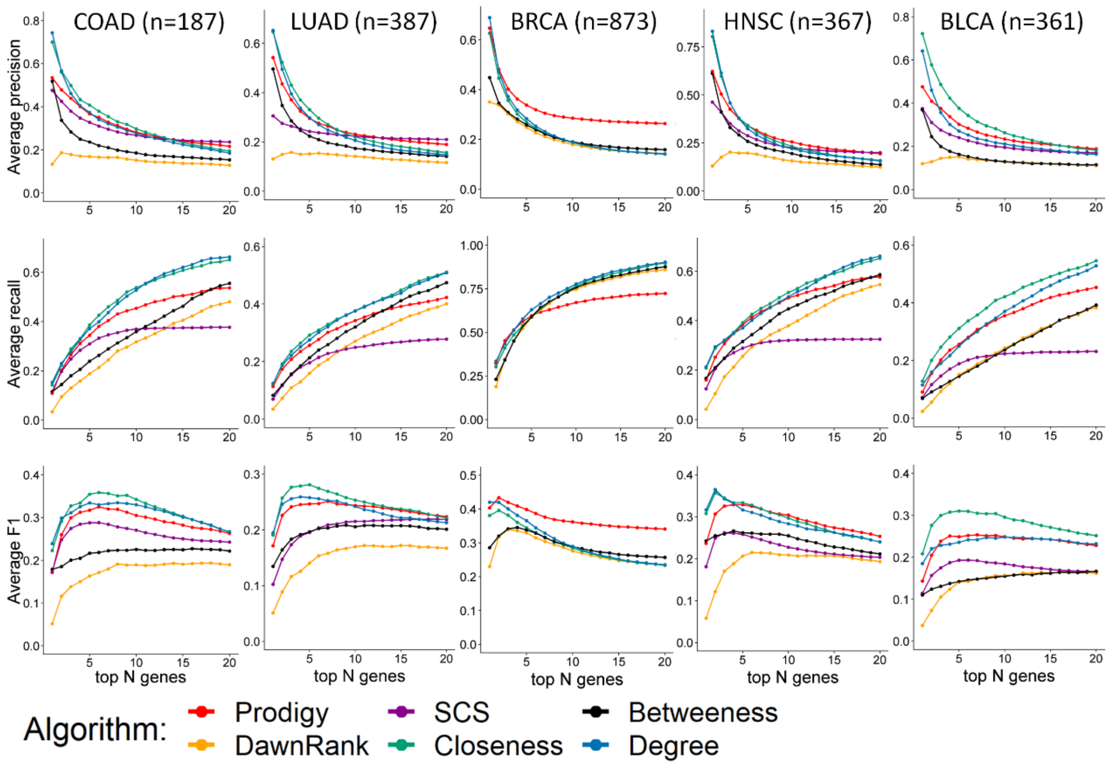


**Supplementary Figure 9**: Performance of the methods on each cohort using NCI pathways. Details are as in Supplementary Figure 7 but here the NCI pathway DB was used.

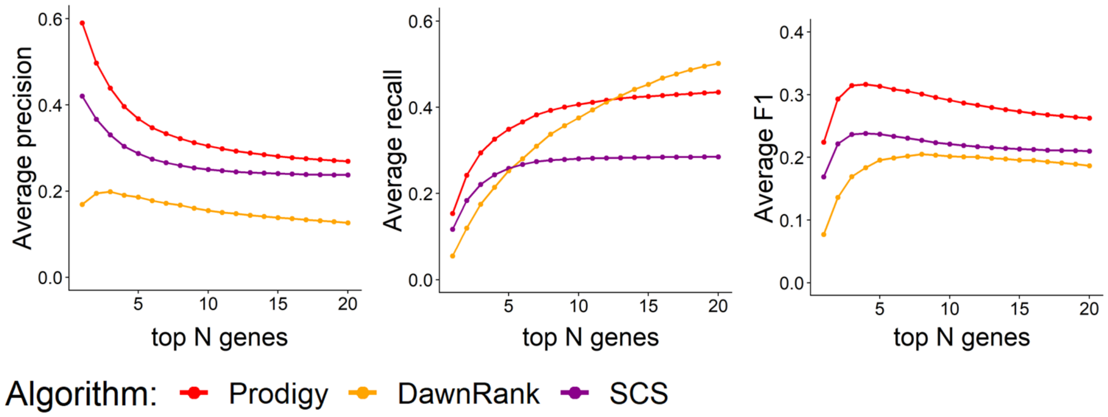


**Supplementary Figure 10**: Average precision, recall and F1 based on the adjusted underlying network from[1,2] (see **Methods**). Results of Prodigy are based on Reactome as pathway DB and for $\alpha = 0.05$. Cohort size: 1804.

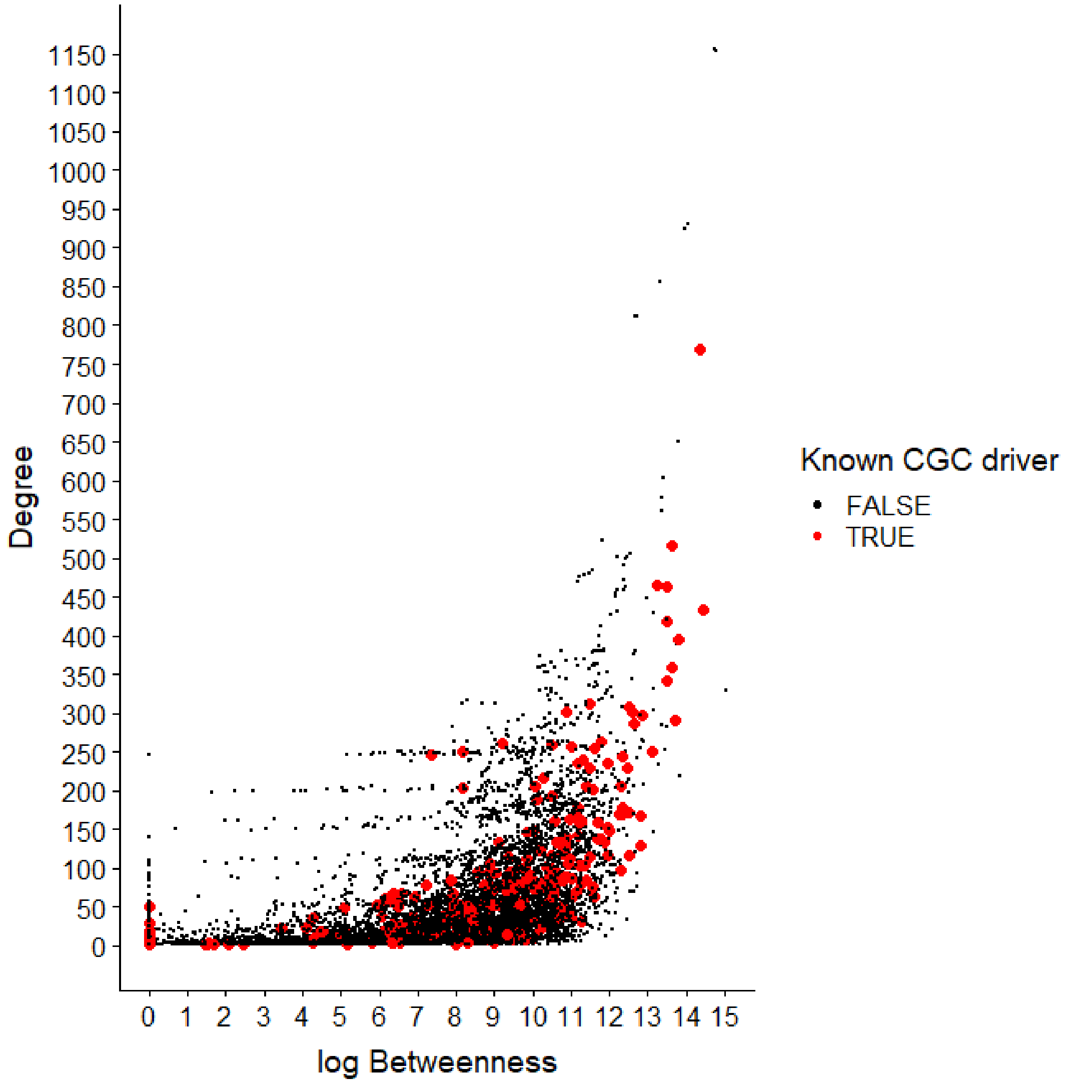


**Supplementary Figure 11:** Centrality measures for all nodes in the global STRING network used in this study. Each point represent a different gene in the network (n = 11,302). The position on the X axis is the log betweenness of the gene and the Y axis is its degree. Known drivers (from CGC[3]) are colored red. Known drivers tend to have higher degrees in the network and higher betweenness values (Wilcoxon rank sum test p-value < 2.2^-16 for both).

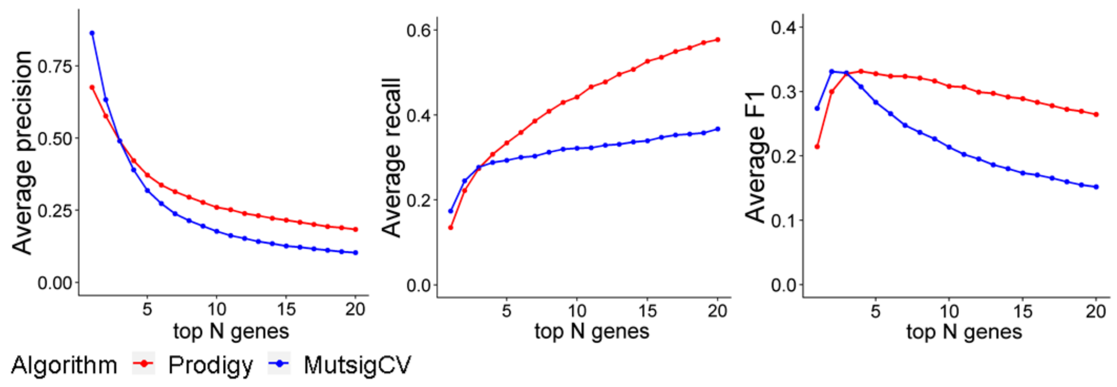


**Supplementary Figure 12**: Results of Prodigy and MutsigCV for a cohort of 178 LUSC patients. Prodigy's results are based on Reactome as pathway DB and for $\alpha = 0.05$.

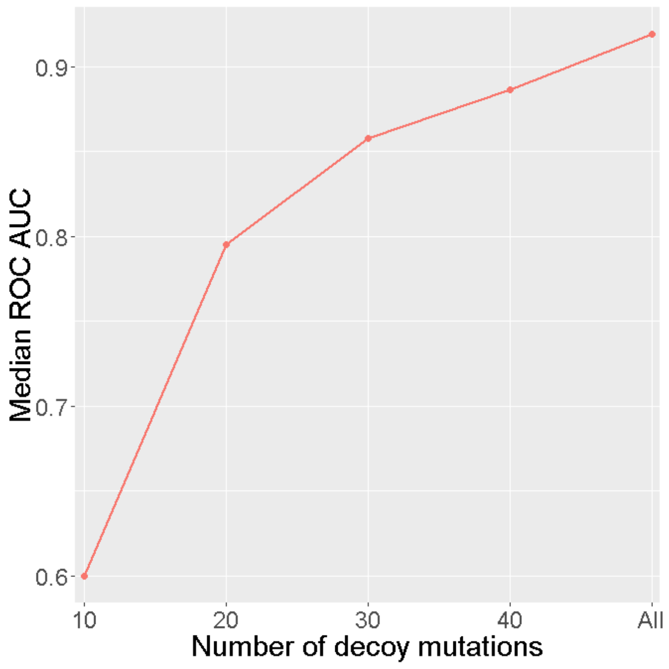


**Supplementary Figure 13**: Discriminating the 10 top genes predicted by Prodigy from a set of decoy mutations. Results are presented as Median AUC across 212 COAD patients. The X axis denotes the number of decoy mutations used.

| Cohort/Number of mutations per mutated gene (%) | 1 | 2 | 3 | >3 |
|---|---|---|---|---|
| COAD | 66817 (91.1) | 5167 (7.04) | 910 (1.24) | 423 (0.57) |
| LUAD | 108946 (82.81) | 16218 (12.32) | 1400 (1.06) | 4990 (3.79) |
| BRCA | 16074 (41.17) | 18224 (46.68) | 4126 (10.56) | 614 (1.57) |
| HNSC | 73194 (95.59) | 2984 (3.89) | 263 (0.34) | 122 (0.15) |
| BLCA | 86066 (94.52) | 4209 (4.62) | 553 (0.6) | 221 (0.24) |

**Supplementary Table 1:** Number of non-silent mutations per mutated gene in different cohorts.

| Pathway database | Reactome | KEGG | NCI |
|---|---|---|---|
| Number of pathways used | 1762 | 285 | 212 |
| Mean number of interactions per pathway (SD) | 1019.6 (3078.9) | 417.6 (849) | 109.6 (107.53) |
| Mean number of genes per pathway (SD) | 58.7 (143.31) | 65.2 (59) | 33.5 (20.2) |

**Supplementary Table 2:** Pathway database statistics.

| Mutated gene | Mutation frequency among patients | Prodigy ranking | MutsigCV overall rank | MutsigCV personalized rank |
|---|---|---|---|---|
| KRAS | 2/178 | 2,3 | 8849 | 49,177 |
| JAK1 | 2/178 | 1,2 | 10520 | 218,183 |
| MAPK1 | 1/178 | 3 | 4451 | 87 |

**Supplementary Table 3**: Personalized identification of driver genes in the LUSC cohort. All genes are known drivers for according to CGC[3].

| Cohort | COAD | BRCA | LUAD | HNSC | BLCA |
|---|---|---|---|---|---|
| Number of observed rarely mutated genes (%) | 24,144 (36.1) | 55,279 (93.8) | 26,682 (42.6) | 44,831 (70.1) | 43,248 (53.9) |
| Number of rare mutations in genes ranked by Prodigy (%) | 403 (21) | 4405 (78.4) | 1610 (37.1) | 2026 (52.9) | 1364 (38.6) |

**Supplementary Table 4**: The number of overall and ranked rarely mutated genes. A gene is rarely mutated in a cohort if it has at least one mutation in < 2% of the patients. For example, since 93.8% of all observed mutated genes in BRCA were mutated in < 2% of patients, the overwhelming majority of mutated genes in BRCA patients are rare. The second row reports the number of rarely mutated genes that were ranked in the top 10 by Prodigy.

| Mutated gene | Cohort | Mutation frequency among cohort's patients | #Patients for whom the gene was highly ranked by Prodigy |
|---|---|---|---|
| SRC | COAD | 1/212 | 1/1 |
| FES | HNSC | 5/502 | 2/5 |
| TSC | HNSC | 6/502 | 2/6 |
| HIF1A | LUAD | 2/487 | 2/2 |
| RAD21 | LUAD | 8/487 | 1/8 |
| ARAF | LUAD | 5/487 | 1/5 |
| EED | LUAD | 6/487 | 1/6 |
| TERT | BLCA | 2/399 | 2/2 |
| RB1 | BRCA | 19/969 | 17/19 |
| CTCF | BRCA | 16/969 | 6/16 |
| CDKN1B | BRCA | 10/969 | 7/10 |
| PBRM1 | BRCA | 6/969 | 1/6 |
| CASP8 | BRCA | 12/969 | 6/12 |

| | | | |
|---|---|---|---|
| EP300 | BRCA | 11/969 | 11/11 |
| ERBB2 | BRCA | 19/969 | 16/19 |
| NOTCH1 | BRCA | 8/969 | 8/8 |
| BRCA2 | BRCA | 15/969 | 2/15 |
| KEAP1 | BRCA | 3/969 | 1/3 |
| AKT1 | BRCA | 2/969 | 2/2 |
| ESR1 | BRCA | 7/969 | 4/7 |
| BAP1 | BRCA | 6/969 | 1/6 |
| SMARCD1 | BRCA | 1/969 | 1/1 |

**Supplementary Table 5**: Examples for personalized identification of rare drivers by Prodigy. All genes are known drivers that are specific for the respective cancer types according to CGC[3] and were ranked in the 10 top genes by Prodigy.
